# Toward standardized behavioral analysis in IntelliCage experiments

**DOI:** 10.64898/2026.09.06.749703

**Authors:** Fabrizio Musacchio, Martin Fuhrmann

## Abstract

Automated home-cage systems measure individual behavior in social groups for days to months. Among these systems, the IntelliCage has become a widely used platform for longitudinal and socially embedded behavioral phenotyping. Yet the analysis layer often remains less standardized than the experiment itself: raw exports, phase definitions, exclusion rules, time alignment, and derived behavioral metrics are transformed by lab-specific scripts that are difficult to audit, compare, or reuse. We present *ic-analysis*, an open-source Python toolkit for standardized, scriptable, and shareable IntelliCage workflows. The toolkit separates user-defined experiment metadata and workflow scripts from a reusable analysis core with modular analysis and plotting functions, allowing users to flexibly assemble experiment-specific pipelines without editing package internals. Due to its modular design, the analysis core can be applied to a broad range of experimental paradigms, including general activity, exploratory, motivational, cognitive, and social readouts, rather than being limited to a single fixed protocol. It aligns biological phase windows across staggered cage runs and exports plots together with quantitative result tables, applied settings, and audit files that support reproducible and FAIR reporting. Here, we demonstrate this flexible design in a realistic synthetic place-learning/place-reversal experiment with two mouse groups and deliberately offset cage starts. The workflow recovered the implanted behavioral differences while preserving the required experimental-time alignment. Group A showed stronger endpoint saccharin preference (81.7 ± 1.7% vs. 27.4 ± 3.4%, *p* = 1.9 × 10^−9^), higher liquid uptake, faster place-learning onset (56.6 ± 8.1 vs. 278.3 ± 33.2 visits), and better reversal performance (64.1 ± 1.3% vs. 25.6 ± 1.7% rewarded correct-corner visits) compared to group B. This demonstration shows how standardized, explicitly defined readouts can turn complex IntelliCage exports into interpretable behavioral profiles while preserving the analysis history needed for inspection and reuse. *ic-analysis* therefore provides both a working analysis scaffold and an extensible, community-friendly route toward IntelliCage workflows that are easier to reproduce, compare, extend, and share.

## Introduction

Behavioral neuroscience increasingly depends on assays that can capture behavior over the same temporal scale on which motivation, learning, social context, circadian state, intervention effects, and disease-related phenotypes unfold. Automated home-cage systems address this need by measuring animals with little experimenter interference and without repeatedly moving them into short, unfamiliar test arenas. The IntelliCage is one of the most widely used systems in this family. It houses RFID-tagged mice in social groups, records visits to four operant corners, and can control access to drinking bottles through nose-poke responses, doors, light cues, and reward contingencies [1, 2]. Over roughly two decades, IntelliCage experiments have been used not only for general activity and exploration, but also for spatial learning, reversal learning, place avoidance, sequencing and patrolling tasks, reward-seeking, taste preference, stress-related behavior, social behavior, pharmacological interventions, and genetic disease models [1, 3, 4] (Table 1).

**Table 1:** Representative IntelliCage behavioral domains and analysis tools motivating standardized, shareable analysis workflows. The publication list maps reported IntelliCage applications to broad behavioral domains and is intended as a landscape overview rather than a formal systematic review.

| Domain | Adaptable paradigms or workflow role | Representative references |
| --- | --- | --- |
| *Behavioral paradigms adaptable to IntelliCage* |  |  |
| General activity and exploration | Free exploration, long-term home-cage activity, circadian activity patterns, new-environment response | [11, 12, 13, 14, 15, 16, 17, 18, 19, 20, 21, 3, 22, 23, 4, 24] |
| Learning, memory, and behavioral flexibility | Place learning, place reversal, place preference and extinction, place avoidance, sequencing, chaining, side learning, and patrolling tasks | [11, 13, 12, 25, 26, 27, 28, 4, 29, 17, 30, 31, 32, 33, 34, 35, 19, 36, 37, 38, 39, 40, 41, 42, 43, 44, 45, 46, 47, 48, 18, 49, 50, 51, 52, 53, 54, 55, 56, 57, 16, 58, 20, 14, 24, 59, 60, 3, 61, 62, 63, 64, 65, 66, 67] |
| Motivation, reward, and decision-making | Taste and sucrose/saccharin preference, reward seeking, alcohol access, conditioned drinking suppression, competition for reward access, impulsivity, and delay discounting | [68, 69, 70, 71, 72, 73, 74, 53, 75, 32, 3, 76, 77, 78, 79, 62, 65, 80, 81] |
| Affective and stress-related behavior | Light response, novel object or odor response, neophobia, cued punishment, chronic unpredictable mild stress, and social stress | [66, 82, 75, 22, 3, 17, 15, 21, 68, 32, 61, 65, 79, 71, 83, 70, 69, 23, 77, 78, 76, 59] |
| Social and group-embedded behavior | Group place learning, social competition, competition for reward access, social modulation of aversive memories, and socially embedded decision-making | [25, 63, 34, 76, 59, 77, 72, 3, 78, 79, 61, 64] |
| Experimental perturbations | Genetic models, pharmacological treatment, longitudinal interventions, disease-model phenotyping, and multimodal studies combined with cellular, histological, or structural analyses | [28, 69, 83, 70, 71, 32, 82, 37, 40, 49, 53, 24, 84, 73, 74, 72, 25, 26, 27, 29, 85, 31, 12, 33, 35, 36, 76, 75, 22, 15, 86, 41, 68, 43, 44, 45, 46, 47, 48, 64, 50, 62, 54, 55, 56, 57, 16, 17, 58, 14, 87, 59, 60, 67, 88, 18, 42, 89, 20, 23] |
| *Analysis tools and workflow concepts* |  |  |
| PyMICE | Open Python library for loading and analyzing IntelliCage data with a strong data-access and scripting layer, leaving experiment-specific behavioral definitions to downstream workflows | [5, 6] |
| IntelliPy | Graphical user interface for common IntelliCage analysis modules, designed to lower the barrier for users without extensive programming experience | [7] |
| IntelliR | R-based standardized pipeline for a defined higher-cognition test battery, including cognition-index outputs across spatial, episodic-like, and working-memory challenges | [8, 9] |
| FlowR | Commercial workflow environment for IntelliCage-related analysis and visualization, not an open, inspectable community code base | [10] |

This breadth is scientifically valuable, but it also exposes a bottleneck. The apparatus records rich event-level data in a comparatively standardized way. The downstream analysis is much less standardized. A single experiment may require decisions about how raw exports are merged, how interrupted runs are concatenated, how biological phases are mapped onto real timestamps, which animals are included, how day/night windows are defined, which visits count as successful, how learning onset is operationalized, and how derived ratios are computed. In many laboratories, these transformations are performed by project-specific scripts or interactive spreadsheets. Such workflows can be effective locally, but they are often difficult to audit, difficult to rerun, and difficult for another laboratory to adapt. The consequence is not merely technical inconvenience. If two IntelliCage studies differ in their handling of phase windows, reward attempts, denominator definitions, or exclusion rules, the resulting behavioral readouts may be less comparable even when the experimental protocols look similar.

The need for automated and transparent analysis has been recognized before. PyMICE provides an open Python library for loading and working with IntelliCage data, with an object-oriented API and support for workflow configuration and validation [5, 6]. IntelliPy wraps such analysis needs in a graphical interface intended to make common IntelliCage workflows accessible to users with little programming experience [7]. More recently, IntelliR presented a standardized R pipeline for automated higher-cognition profiling across defined IntelliCage challenge designs, including spatial, episodic-like, and working-memory tasks with reversals [8, 9]. A proprietary option, FlowR, is also available as part of IntelliCage-related commercial workflows [10]. These tools demonstrate that the field has moved beyond manual extraction.

At the same time, they occupy different positions in the analysis landscape: general data-access libraries still leave many behavioral definitions to the user, graphical interfaces can trade flexibility for ease of use, and challenge-specific pipelines can be highly standardized while being less immediately general to other experimental layouts. Table 1 places this tool landscape next to representative IntelliCage behavioral domains. Its purpose is to show why analysis standardization matters now: IntelliCage protocols are no longer a narrow niche of spatial learning, and analysis tools are no longer absent. The unmet need is a workflow layer that is open, scriptable, modular, transparent about how each behavioral metric is defined, and easy to extend to new experiment classes without forcing users to edit core package code.

### Box 1

**Design goals for standardized IntelliCage analysis**

1. **Separate biological phases from export blocks**. Raw IntelliCage exports may reflect interruptions, restarts, or convenient export chunks. Analysis should instead use explicit experiment- and subject-level phase windows.
2. **Keep the user workflow scriptable and readable**. Users should define experiment metadata, subject metadata, and the analysis pipeline in a transparent script without editing the package core.
3. **Define behavioral metrics explicitly**. Each analysis metric should be operationally and, where applicable, mathematically defined, including the events, windows, transformations, and summary rules from which it is computed.
4. **Pair every figure with reusable data**. Plots should be generated together with CSV tables, machine-readable settings, and audit files rather than as isolated visual products.
5. **Support modular rather than monolithic analysis**. Users should be able to run only the analyses appropriate for their protocol and extend the package with new readouts.
6. **Favor open and community-extensible standards**. The analysis layer should be open-source, versionable, and compatible with FAIR-oriented sharing of data, code, and results.

This motivation parallels broader concerns in animal behavior research. Automated systems can reduce some sources of experimenter interference and enable richer longitudinal measurements, but automation alone does not guarantee reproducibility [90]. Reporting standards such as ARRIVE and data stewardship principles such as FAIR emphasize that studies should be described, structured, and shared in ways that make their assumptions and outputs findable, accessible, interoperable, and reusable [91, 92]. For IntelliCage experiments, this means that the analysis should not end in a figure exported from a local notebook. It should also produce the tables, settings, phase definitions, and audit trail needed to understand how that figure was generated.

Together, these considerations motivate the design goals summarized in Box 1. A standardized workflow first has to separate biological experiment phases from technical export blocks, because export folders often reflect recording interruptions or convenient file handling rather than the actual behavioral protocol (goal 1). It should keep experiment definitions readable at the user-script level, so that animal inclusion, group assignment, phase timing, and selected analyses remain visible to the researcher (goal 2). The behavioral quantities themselves must be explicit enough that another laboratory can understand which events, windows, transformations, and summaries entered a result (goal 3). Figures should be coupled to reusable result tables and run settings rather than existing as isolated images (goal 4). The analysis layer should remain modular, so that users can assemble the functions needed for a given experiment and extend the package to new readouts (goal 5). Finally, the whole workflow should be open and versionable, which makes it compatible with community review, reuse, and FAIR-oriented sharing (goal 6).

To address this design brief, we developed *ic-analysis*, an open-source Python toolkit for standardized IntelliCage workflows. The toolkit is not intended as another closed analysis recipe for a single IntelliCage protocol. Instead, it separates user-defined experiment metadata and workflow choices from a reusable analysis core, maps raw export folders into an aligned experiment timeline, computes behaviorally defined metrics from event-level visits and nosepokes, and writes reproducible result packages. Figure 1 summarizes the conceptual workflow. IntelliCage experiments unfold as biological phases but may be exported as arbitrary data blocks. The user script defines the experiment, phases, subjects, and desired analyses. The package core loads, aligns, and prepares the event tables, and each analysis run produces plots, CSV tables, settings, and audit files that can be inspected or shared. By turning phase definitions, metric construction, plotting, and output provenance into reusable but adaptable workflow components, *ic-analysis* is intended to serve as a shared analysis layer through which IntelliCage laboratories can compare results more directly, adapt assays without rewriting core code, and extend standardized behavioral readouts as new experimental paradigms emerge.

**Figure 1:**
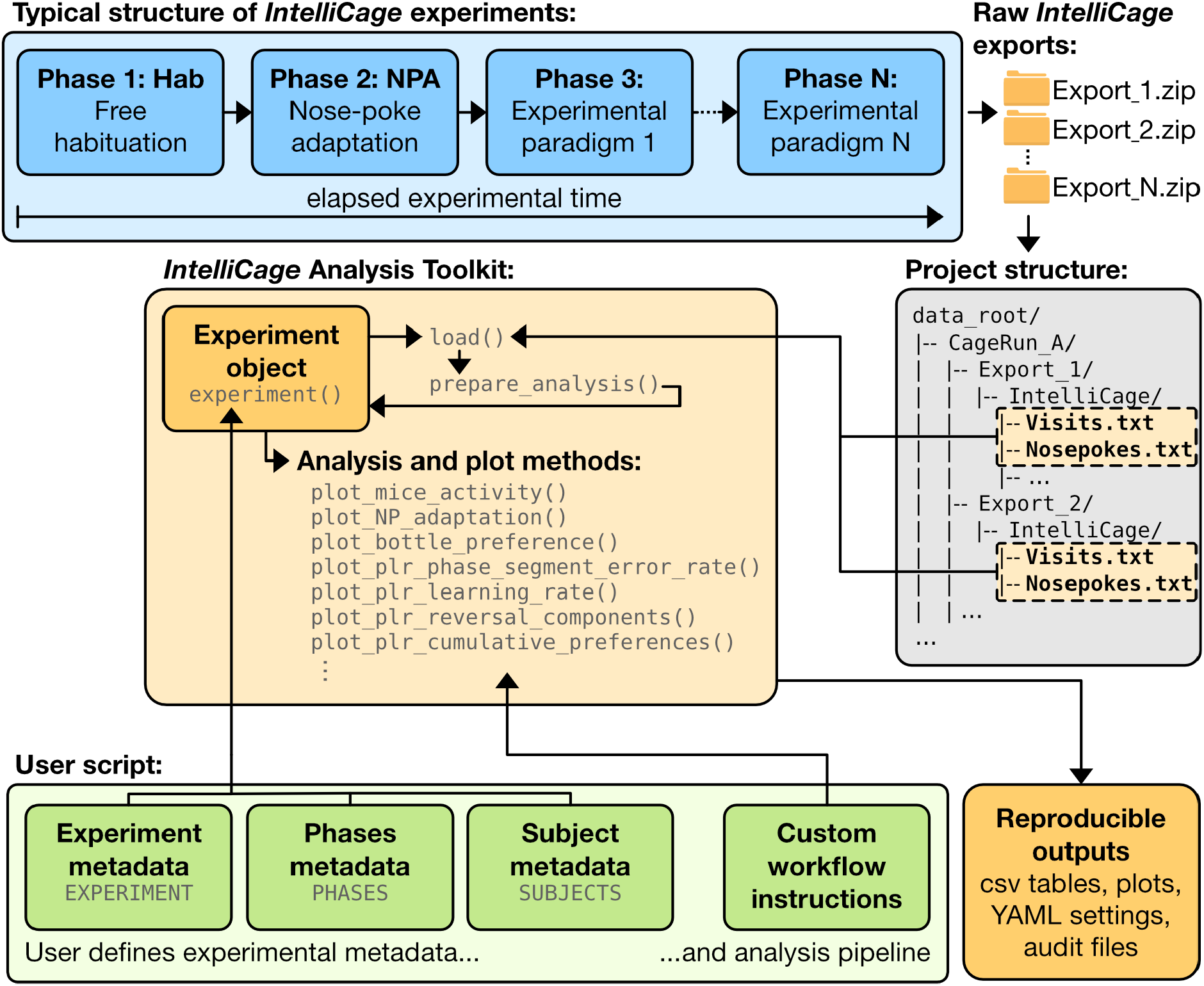
Conceptual structure of a standardized IntelliCage analysis workflow. IntelliCage experiments are planned as biological phases, but raw data often arrive as export blocks that may or may not coincide with phase transitions. In *ic-analysis*, the user script defines experiment metadata, phase metadata, subject metadata, and a custom sequence of analysis calls. The package core keeps loading and preparation separate from modular analysis and plotting methods. Each run writes reproducible outputs, including CSV tables, plots, YAML settings, and audit files.

## Results

### A synthetic place-learning/place-reversal experiment as a behavioral test case

To demonstrate the analysis workflow, we generated a synthetic IntelliCage dataset with two experimental groups and a four-phase place-learning/place-reversal (PLR) protocol outlined in Figure 2a. The dataset mimics standard IntelliCage exports by storing subject-resolved visit events, nose-poke events, and lick records in tabular files. This protocol is used here as a demanding worked example, not as the boundary of the toolkit. The same user-script and reusable-core structure is intended to support general activity, exploratory behavior, drinking and reward-related assays, cognitive tasks, and social or group-embedded experiments, provided that the relevant behavioral events and phase windows are defined explicitly. The synthetic PLR protocol was designed to resemble a common appetitively motivated IntelliCage workflow. During free habituation (Hab), all corners were accessible. During nose-poke adaptation (NPA), access required one nose poke. During place learning (PL), mice were assigned to individual rewarded corners, while during place reversal (PR), each mouse was reassigned to a new corner, forcing a change in reward location. Throughout the experiment, each corner contained one bottle of sweetened water and one bottle of plain water in the same side arrangement. This configuration served two purposes: sweetened water provided the appetitive liquid used for reward-related behavior, and the two-bottle setup allowed the analysis to separate total liquid uptake from relative saccharin preference. The synthetic dataset was designed to implant a behavioral difference between the two groups. Group A was designed with stronger place learning and stronger sweetened-water engagement, whereas Group B was designed with weaker place learning and weaker sweetened-water engagement. The synthetic dataset was generated to be realistic, but it is not intended to represent a specific biological experiment or to be interpreted as a real behavioral outcome.

**Figure 2:**
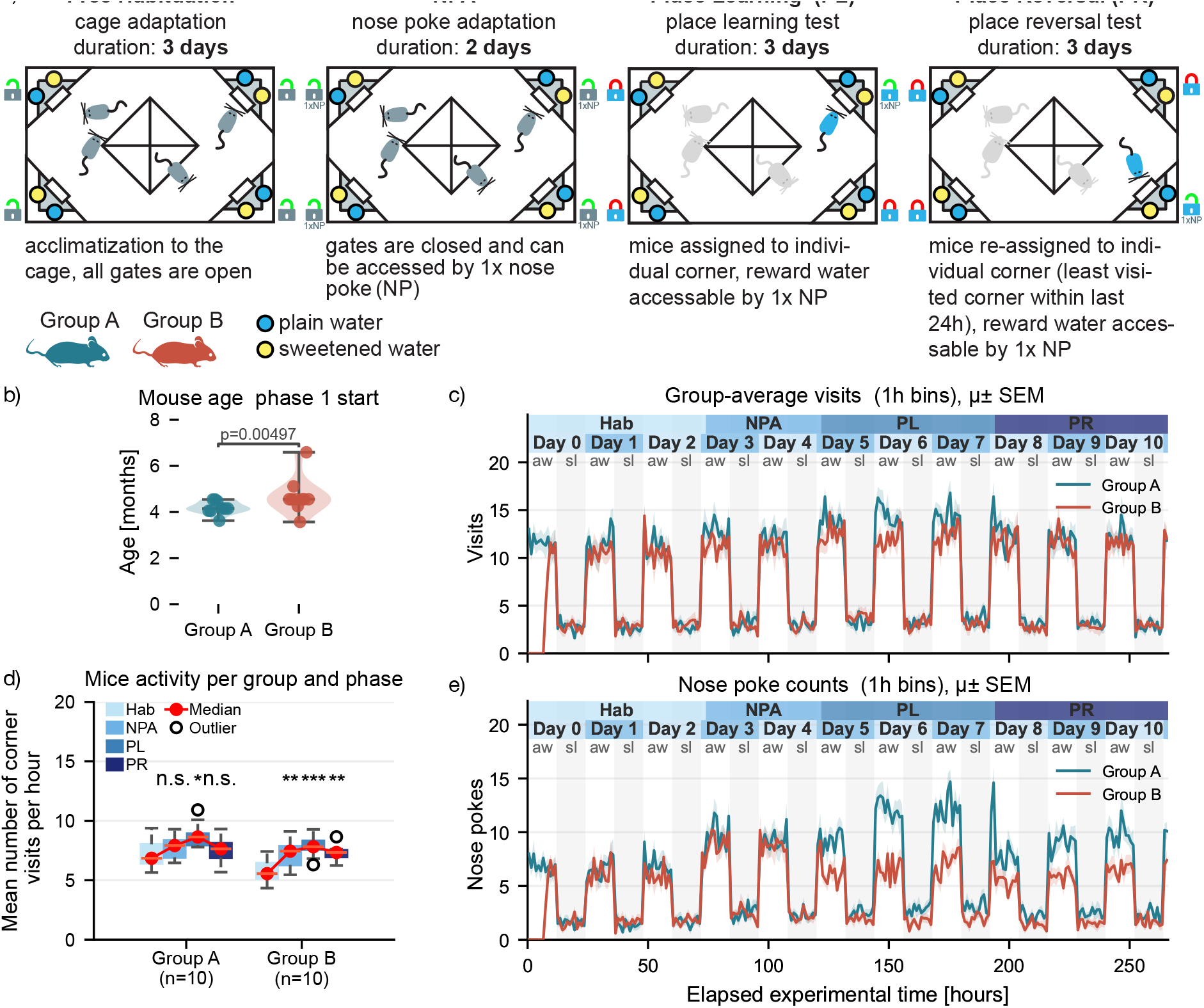
Protocol overview, metadata, activity, and nose-poke adaptation in the synthetic PLR dataset. **a**, Four-phase place-learning/place-reversal protocol with free habituation, NPA, PL, and PR. Yellow and blue bottle symbols indicate sweetened and plain water. **b**, Mouse age at phase 1 start. **c**, Group-average all-corner visit activity in 1 h bins. **d**, Phase-wise per-mouse activity summary. **e**, Nose-poke counts in 1 h bins. Lines and shaded bands show group means ± SEM where applicable.

The dataset was deliberately not perfectly aligned at the raw timestamp level. Group B was simulated as a separate cage run starting 12 days later than Group A, and its first detected phase-1 entries occurred 7.5 h after the beginning of the mouse day. This created a stress test for the package: a correct analysis should align behavior to biological experiment time and phase windows, not to absolute calendar dates or export folder starts. The resulting overview plot shown in Figure 2c confirmed that group-level activity traces could be represented on a shared elapsed experimental timeline while preserving the delayed phase-1 entry of Group B. The two groups also differed modestly in age at phase 1 start (Group A: 4.19 ± 0.09 months; Group B: 4.68 ± 0.24 months; *p* = 0.00497), illustrating how subject metadata can be inspected before interpreting behavioral outcomes (Figure 2b).

General visit activity provided an immediate quality-control view of the experiment. Both groups showed strong day/night modulation across the experiment, with high activity during the awake phase and lower activity during the sleep phase (Figure 2c). Phase-wise activity summaries indicated that Group A increased from 7.26 ± 0.37 visits per hour in habituation to 8.78 ± 0.32 visits per hour during PL, whereas Group B increased from 5.75 ± 0.29 visits per hour in habituation to 7.82 ± 0.26 visits per hour during PL (Figure 2d). These changes are not the main biological endpoint in this demonstration, but they are important controls: differences in learning and drinking should be interpreted against the background of overall cage engagement.

The NPA readout shown in Figure 2e served a second quality-control role. Both groups acquired substantial nose-poke activity during the adaptation phase, with mean nose-poke counts of 6.00 ± 0.25 events per hour in Group A and 5.60 ± 0.27 events per hour in Group B. Thus, before the formal place-learning phase began, the workflow showed that both groups interacted with the operant doors at comparable levels. This helps separate later differences in drinking preference and learning performance from a simple failure to acquire nose-poke access.

### Standardized drinking metrics separate total uptake from saccharin preference

The same event tables were then used to quantify drinking behavior (Figure 3). The distinction between absolute uptake and relative preference is behaviorally important. A mouse can show a high preference for a sweetened bottle while drinking little overall, or can drink substantially while showing weak preference. The toolkit therefore reports both total consumption and side- or bottle-specific ratios.

**Figure 3:**
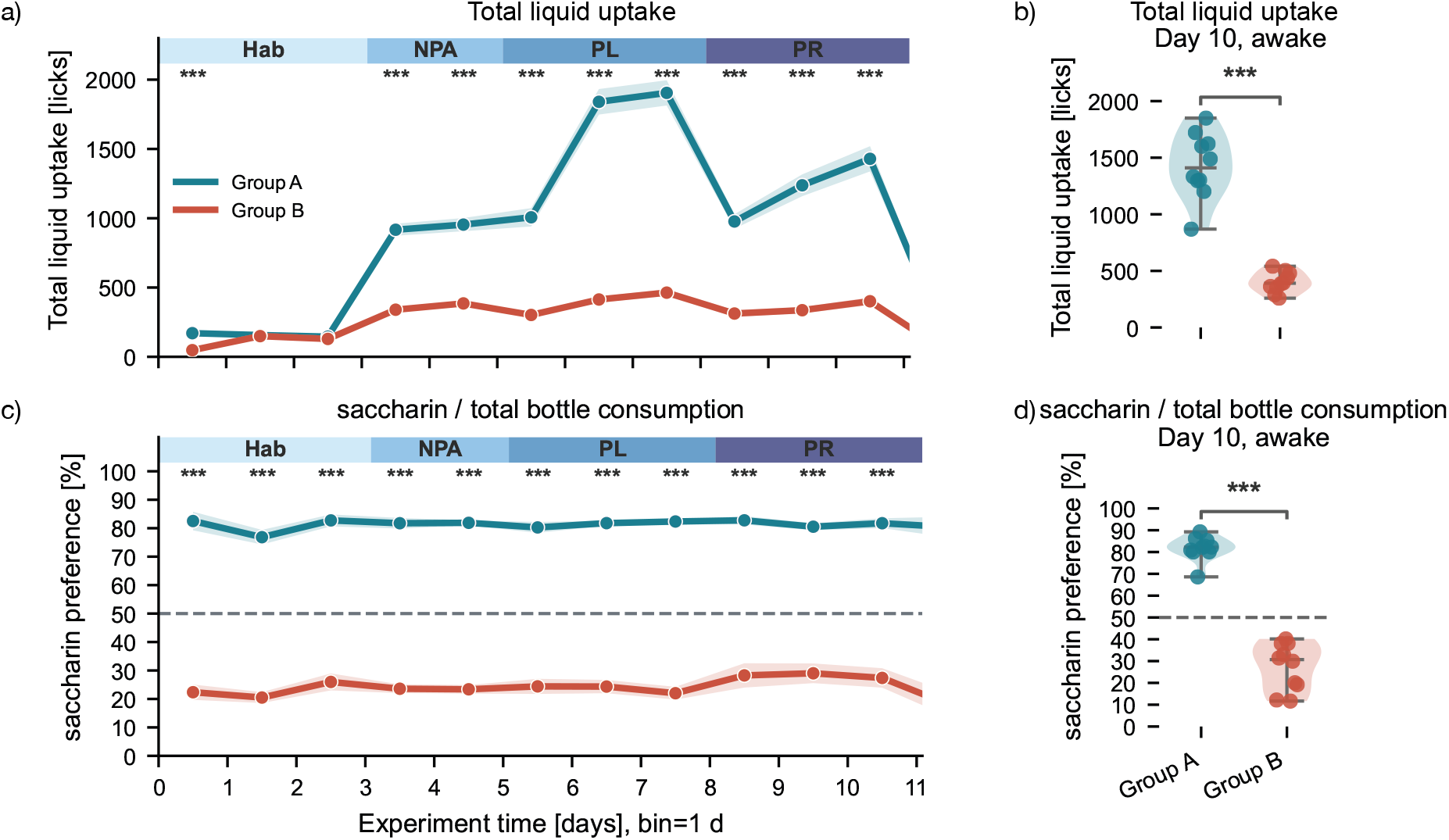
Drinking behavior in the synthetic PLR dataset. **a**, Total liquid uptake across the experiment in 24 h bins. **b**, Total liquid uptake on experimental day 10 during the awake phase. **c**, Saccharin preference, defined as saccharin-bottle consumption divided by total bottle consumption. **d**, Saccharin preference on experimental day 10 during the awake phase. Dashed horizontal lines indicate 50% preference where applicable.

Across the full experiment, Group A showed higher total liquid uptake than Group B (Figure 3a). On experimental day 10 during the awake phase, Group A produced 1428.8 ± 90.8 licks compared with 400.9 ± 28.8 licks in Group B (Welch test, *p* = 4.1 × 10^−7^; Figure 3b). This difference reflects the synthetic design, in which Group A was assigned stronger sweetened-water engagement. Relative saccharin preference showed an even clearer group separation. Group A remained strongly biased toward the saccharin bottle across the experiment, whereas Group B remained biased toward the alternative bottle or showed only weak saccharin preference (Figure 3c). On day 10 during the awake phase, Group A reached 81.7 ± 1.7% saccharin preference, while Group B reached 27.4 ± 3.4% (Welch test, *p* = 1.9 × 10^−9^; Figure 3d). These paired views demonstrate why standardized analysis should keep raw consumption and relative preference separate: the former quantifies total fluid interaction, whereas the latter quantifies choice bias among available bottles.

### Place learning and reversal readouts reveal acquisition, errors, and perseveration

We next analyzed the core PL and PR behavior as rewarded correct-corner visit rates and their complementary error rates (Figure 4). During PL, Group A increased from 40.1 ± 1.3% rewarded correct-corner visits on phase day 1 awake to 74.1 ± 1.9% on phase day 3 awake. Group B improved more slowly, from 16.8 ± 0.9% to 27.5 ± 1.2% over the same period. Thus, Group A moved from above chance to a strong correct-corner bias, whereas Group B remained close to chance or only moderately above it (Figure 4a,c). The endpoint comparison on PL day 3 awake was highly separated (*p* = 1.1 × 10^−13^).

**Figure 4:**
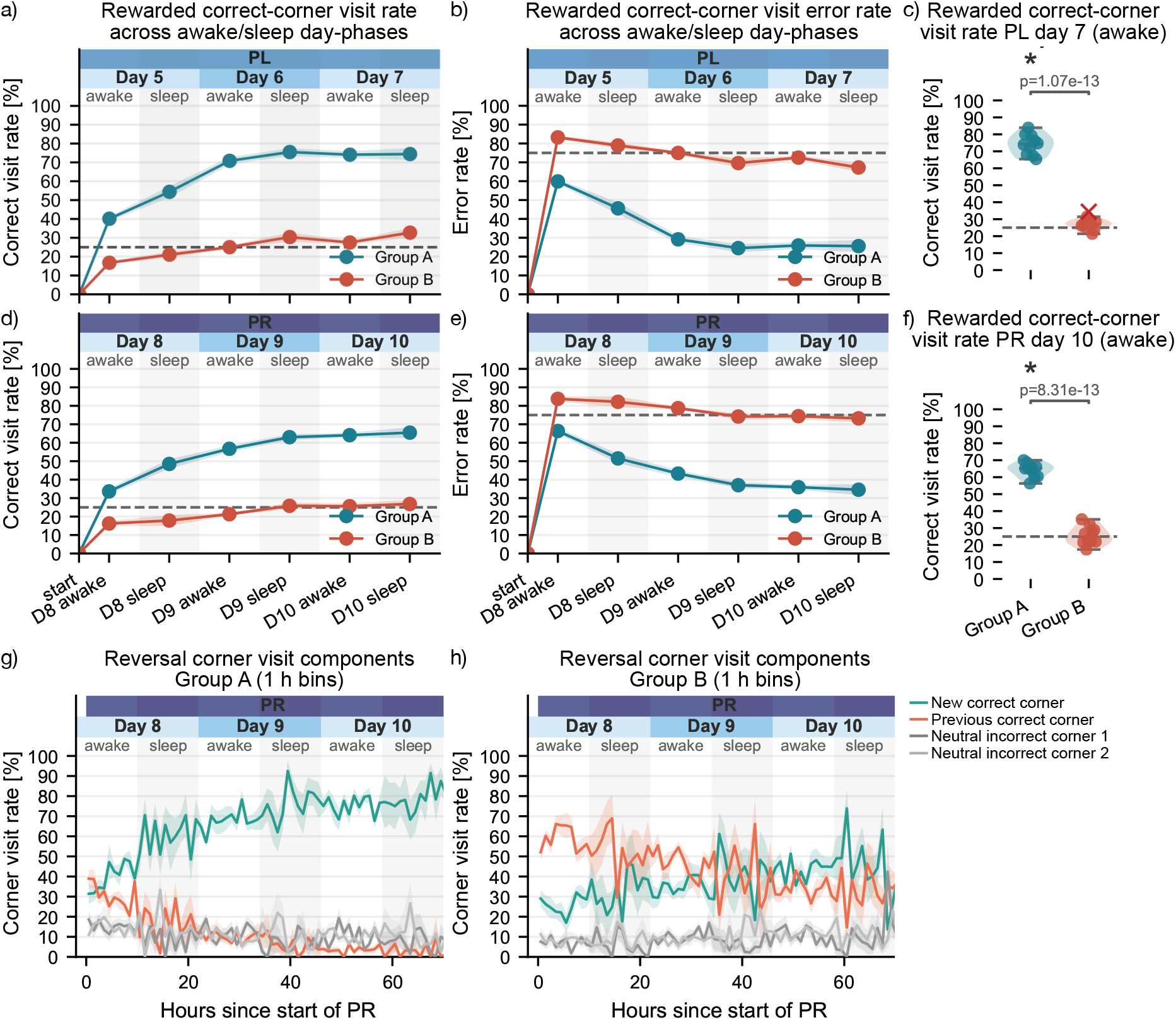
Place learning, place reversal, and reversal components. **a**, Rewarded correct-corner visit rate across PL awake/sleep segments. **b**, Complementary rewarded correct-corner visit error rate during PL. **c**, PL endpoint on phase day 3 awake. **d**, Rewarded correct-corner visit rate across PR awake/sleep segments. **e**, Complementary rewarded correct-corner visit error rate during PR. **f**, PR endpoint on phase day 3 awake. **g**,**h**, PR corner-role decomposition for Group A and Group B, separating new correct, previous correct, and neutral incorrect corners. Dashed horizontal lines indicate the four-corner chance level for correct-corner analyses.

The error-rate view recasts the same denominator as incorrect or unrewarded choices. In PL, Group A reduced its rewarded correct-corner error rate to 25.9 ± 1.9% by day 3 awake, whereas Group B remained at 72.5 ± 1.2% (Figure 4b). Showing the success rate and error rate together is useful because the success curve emphasizes acquisition, whereas the error curve emphasizes the residual behavioral cost of wrong choices.

The reversal phase posed a harder flexibility problem because the previously correct corner could continue to attract visits. Group A again improved across PR, reaching 64.1 ± 1.3% rewarded correct-corner visits on PR day 3 awake. Group B reached only 25.6 ± 1.7% (*p* = 8.3 × 10^−13^; Figure 4d,f). The corresponding PR error rates were 35.9 ± 1.3% for Group A and 74.4 ± 1.7% for Group B on day 3 awake (Figure 4e).

Reversal-component plots decomposed this group difference into behaviorally interpretable corner roles. During the first 6 h of PR, Group A distributed visits between the new correct corner (36.9%) and the previously correct corner (32.5%), then shifted strongly toward the new correct corner by the final 6 h (83.5%). Group B began PR with a dominant previous-corner bias (60.9% previous vs. 23.5% new in the first 6 h) and still showed incomplete switching by the final 6 h (38.4% new, 32.9% previous). This analysis distinguishes poor reversal performance caused by random exploration from poor reversal performance caused by perseveration at the formerly rewarded location.

### Experience-based analyses distinguish exposure from elapsed time

Elapsed time is not always the best axis for learning. In IntelliCage experiments, mice self-initiate visits, and two animals can experience different numbers of trials within the same number of hours. The toolkit therefore includes experience-based learning curves that plot success probability by visit number rather than clock time (Figure 5). In PL, Group A rose rapidly above chance on the experience axis and reached 70.0 ± 3.4% rewarded correct-corner success by approximately 250 visits. Group B reached only 26.5 ± 3.6% at the same experience level and remained on the same low throughout the experiment (Figure 5a). Learning onset was defined as the first rolling visit window from which three consecutive windows exceeded the configured success threshold. By this definition, all Group A mice reached a PL onset, at 56.6 ± 8.1 visits. In Group B, one mouse did not reach onset, and the remaining nine mice did so much later, at 278.3 ± 33.2 visits (*p* = 0.00028; Figure 5b).

**Figure 5:**
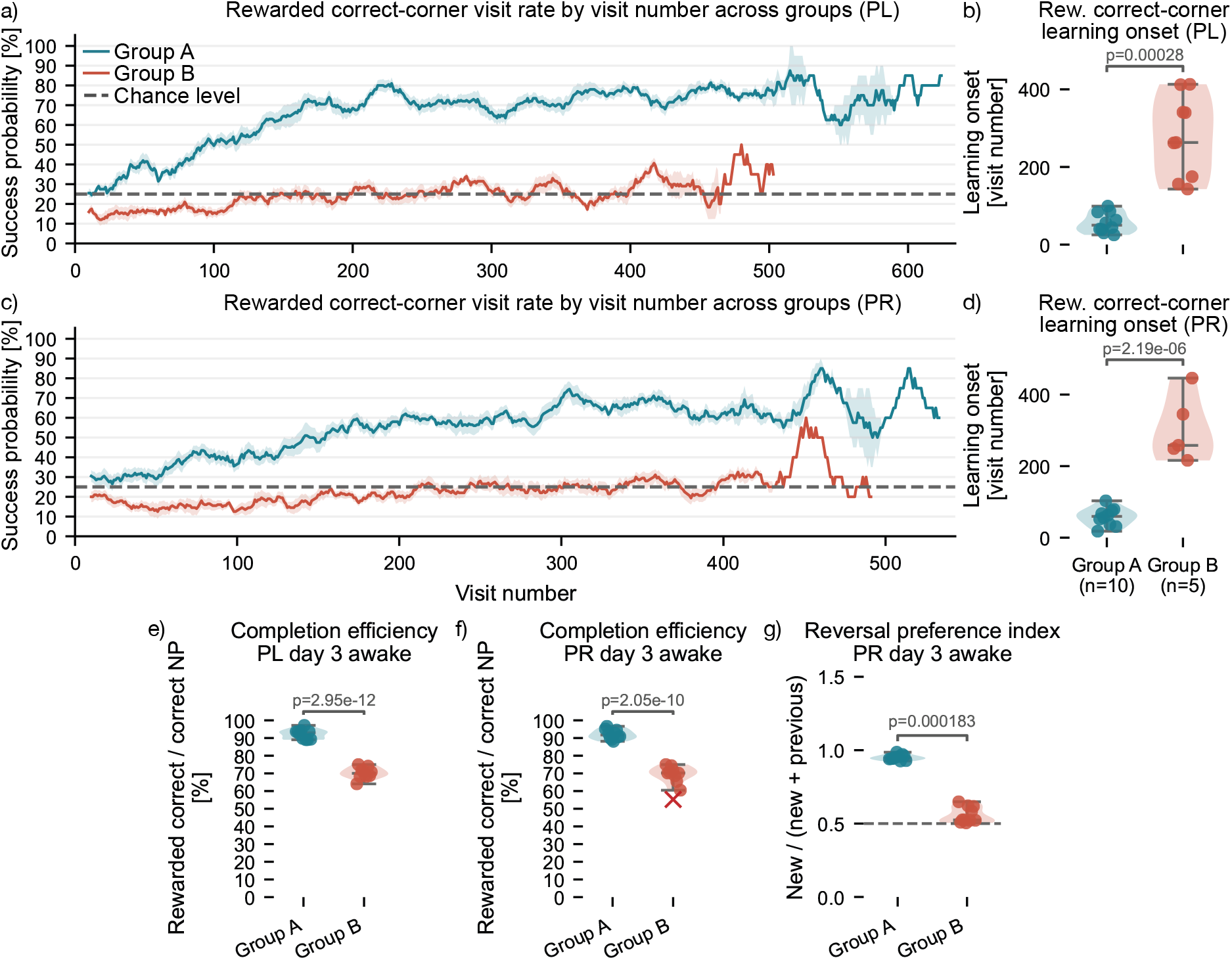
Experience-based learning and derived endpoint ratios. **a**,**c**, Rewarded correct-corner visit rate by visit number during PL and PR. **b**,**d**, Learning onset in visit number for PL and PR. Missing Group B PR onsets indicate mice that did not cross the configured criterion. **e**,**f**, Completion efficiency on phase day 3 awake, defined as rewarded correct-corner visits divided by correct-corner nose-poke visits. **g**, Reversal preference index on PR day 3 awake, defined as new-correct-corner visits divided by visits to the new or previous correct corner.

The same experience-based view showed weaker and less complete learning during PR. All Group A mice crossed the PR onset criterion at 58.1 ± 7.9 visits. Only five of ten Group B mice crossed the criterion, with a mean onset of 302.8 ± 29.5 visits among responders (*p* = 2.2 × 10^−6^; Figure 5c,d). The reduced Group B sample size in the onset plot is therefore not a plotting artifact: mice without an onset are not assigned an arbitrary late value but remain missing for onset statistics.

Derived endpoint ratios provided additional behavioral interpretation. Completion efficiency asked how often a correct-corner nose poke was followed by licking, thereby separating correct approach from successful reward completion. On PL day 3 awake, Group A reached 92.2 ± 0.9% completion efficiency compared with 70.2 ± 1.0% in Group B (*p* = 3.0 × 10^−12^; Figure 5e). On PR day 3 awake, the values were 92.1 ± 0.9% and 68.2 ± 2.0% (*p* = 2.1 × 10^−10^; Figure 5f). Finally, the reversal preference index quantified how strongly PR visits to the two historically relevant corners favored the new PR target over the previous PL target. By PR day 3 awake, Group A approached exclusive preference for the new target (0.951 ± 0.006), whereas Group B remained closer to an even split (0.558 ± 0.017; *p* = 0.000183; Figure 5g).

## Discussion

The central premise of this work is that IntelliCage standardization should not stop at the hardware, the protocol, or the raw export. It must include the analysis layer. The IntelliCage field already spans general activity, circadian patterns, spatial learning, reversal, reward-seeking, taste preference, social behavior, stress-related phenotypes, intervention studies, and genetic disease models. This breadth makes the system powerful, but it also makes post hoc analysis choices more consequential. A “correct-corner rate” is not a universal object unless the denominator is defined. A “learning onset” is not comparable unless the window size, threshold, and crossing rule are defined. A phase-specific endpoint is not interpretable unless the mapping from raw timestamps to biological phase windows is explicit.

*ic-analysis* addresses this problem by proposing a middle position between a general data-access library and a fixed challenge-specific pipeline. Like PyMICE, it is open and scriptable in Python [5]. Unlike a pure data-access layer, it provides ready-to-run behavioral analyses and plots with explicit output tables. Like IntelliPy, it aims to make common analysis operations easier for users [7]. Unlike a GUI-first approach, the user-facing object is a versionable script that can be inspected, copied, diffed, and shared. Like IntelliR, it embraces standardized analysis of IntelliCage cognition paradigms [8]. Its main difference is that standardization is implemented through modular user-level workflows rather than through a single fixed challenge design. Users define phases, subjects, group assignments, and selected methods for the experiment at hand, while the package core handles common loading, alignment, metric computation, plotting, and provenance. Thus, the PLR example shown here should be read as one implementation of the design rather than as its scope limit. The same architecture can be extended to other activity, exploration, motivation, cognition, and social-behavior paradigms by adding behaviorally explicit metrics.

The synthetic dataset was designed to expose this problem in a controlled setting. It included a 12-day difference between cage runs and a 7.5 h delayed phase-1 entry in one group. A brittle workflow tied to calendar dates, export-folder names, or the first observed event in every selected plot would misalign activity with day/night state or phase boundaries. The workflow used here instead relies on subject-level phase windows and elapsed experimental time. This allows standard activity, NPA, drinking, PL/PR, reversal, and experience analyses to be interpreted in one coherent timeline.

The behavioral phenotype recovered from the synthetic data is intentionally strong, but it is useful precisely because it exercises multiple levels of analysis. The synthetic group A showed higher drinking motivation, strong saccharin preference, rapid place learning, successful reversal, high completion efficiency, and a shift away from the previously correct corner. The synthetic group B showed lower liquid uptake, low saccharin preference, slower or missing learning onsets, weaker reversal, lower completion efficiency, and persistent previous-corner attraction. Taken together, these readouts form a coherent behavioral and cognitive profile rather than a set of disconnected endpoint measures. The value of the standardized workflow is that this profile is assembled from explicit event-level definitions on a shared experimental timeline, making the interpretation easier to inspect, reproduce, and extend.

Several limitations follow from the current scope. First, the demonstration dataset is synthetic. It is useful for testing alignment, output consistency, and expected metric behavior, but it is not evidence for a biological group difference. The package should therefore be validated further on independent real IntelliCage datasets and across laboratories. Second, although the current release already supports general activity, liquid-intake surveillance, nose-poke adaptation, bottle preference, place learning, and place reversal workflows, many IntelliCage paradigms in Table 1 will require additional experiment-specific readouts. The toolkit is designed for that extension, but the extension itself still needs careful behavioral definition. Third, the implemented statistical tests are intended as transparent defaults, not replacements for preregistered inferential models. Longitudinal IntelliCage data can involve repeated measures, cage effects, sex effects, circadian structure, and missingness patterns that may require mixed models or custom statistics.

For standardization to become useful in practice, analysis conventions must be easy to adopt without forcing experiments into a single fixed protocol. *ic-analysis* supports this by giving users a transparent script-level workflow for current analyses and an extensible structure in which new readouts can be added with explicit behavioral definitions and reproducible outputs. Because of its open-source and modular structure, it is also open to community-contributed extensions, enabling its long-term evolution alongside the growing diversity of IntelliCage experiments. By making these workflows easier to assemble, inspect, rerun, share, and extend, the toolkit supports FAIR-aligned community standards for longitudinal, socially embedded home-cage behavior.

## Methods

### Toolkit architecture and workflow design

Figure 1 summarizes the separation between biological experiment structure, raw data export, user-level configuration, and package-internal analysis. The upper blue panel represents the intended IntelliCage experiment as a sequence of biological phases along elapsed experimental time. The raw output shown on the right is organized differently. It arrives as one or more export archives, which are unpacked into cage-run and export-block directories containing tabular files such as `Visits.txt` and `Nosepokes.txt`. These export blocks are technical data containers. They may reflect interruptions, restarts, or convenient export intervals and are therefore not treated as phase definitions. This separation addresses design goal 1 in Box 1.

The center of the toolkit, indicated by the orange core in Figure 1, is the experiment object. It receives three user-defined metadata structures. Experiment metadata (`EXPERIMENT`) define paths, group names, colors, and the mouse day. Phase metadata (`PHASES`) define phase labels, display colors, and scheduled phase starts. Subject metadata (`SUBJECTS`) define inclusion, group assignment, sex, date of birth, readable IDs, phase-specific time windows, and place-learning corner assignments. Keeping these scientific choices in the user script makes the analysis readable and directly supports design goal 2 in Box 1.

The core methods `load()` and `prepare_analysis()` connect the raw exports to the biological analysis timeline. During loading, the package scans cage-run folders for all export blocks and concatenates the raw visit and nose-poke tables. During preparation, raw event tables are merged with subject metadata, assigned to biological phase windows, aligned to elapsed experimental time, annotated with day/night state, and enriched with event flags used by downstream metrics. The most important step is phase alignment across cage runs. It allows animals recorded on different calendar dates, or entering the cage at different times within the mouse day, to be analyzed on the same biological experiment axis.

The lower green panel in Figure 1 represents the user script. The user defines the `EXPERIMENT`, `PHASES`, and `SUBJECTS` structures and then assembles a custom analysis workflow by calling the desired methods, for example activity, bottle-preference, NPA, PLR, reversal, or experience-based analyses. This design makes the workflow modular rather than monolithic. Users can run only the analyses appropriate for their protocol, and new readouts can be added without changing the loading and alignment machinery. This directly supports design goal 5 in Box 1.

Each analysis call writes reproducible outputs, shown in the lower right orange box of Figure 1. These include plots, TSV/CSV-compatible tables, applied settings, metadata snapshots, phase manifests, and audit-oriented intermediate files. The same run therefore produces both publication-oriented figures and the machine-readable data needed to inspect or rerun them. This output design implements the figure/table coupling and FAIR-oriented sharing goals described in design goals 4 and 6 of Box 1.

In the following sections, we describe the key concepts and mathematical formalism underlying the experiment time, behavioral metric definitions, and export summaries.

### Experiment time, export blocks, and biological phase windows

Let *m* index mice and *i* index events. Each event has an absolute timestamp *t*_*i*_. For each mouse and biological phase *p*, the user may define a subject-specific phase window

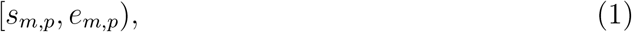

where *s*_*m,p*_ and *e*_*m,p*_ are real timestamps. An event is assigned to phase *p* if

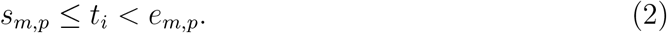

The corresponding phase-relative elapsed time is

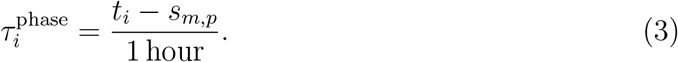

The experiment-relative elapsed time is computed relative to the run-specific experiment-day anchor *a*_*m*_,

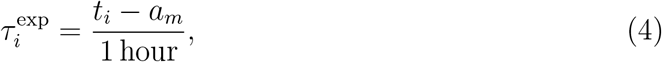

with optional schedule-based alignment when subject-specific phase windows are not supplied. This phase-window assignment addresses design goal 1 in Box 1 by separating biological protocol phases from technical export blocks.

### Visit-level event definitions

The prepared visit table stores event-level indicators used by downstream analyses. To make behavioral metrics explicit as required by design goal 3 in Box 1, these indicators are defined from visit, nose-poke, lick, and corner-assignment information before downstream aggregation. Let *c*_*i*_ be the visited corner, *n*_*i*_ indicate that the visit contained at least one nose-poke event, and *ℓ*_*i*_ indicate lick-positive reward access. For place-learning and reversal phases, each mouse has a phase-specific assigned corner *a*_*m,p*_. The basic correct-corner indicator is

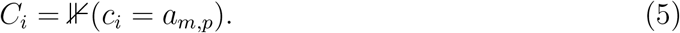

The correct-nose-poke indicator is

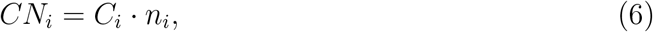

and the rewarded correct-corner visit indicator is

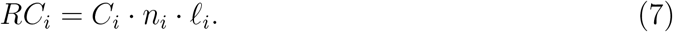

For place reversal (PR), the previously correct corner is the place-learning (PL) assigned corner. The previous-corner indicator is therefore

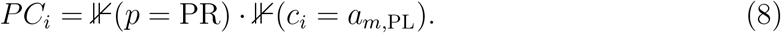

Neutral incorrect corners are corners that are neither the new PR corner nor the previous PL corner. During NPA, lick-positive nose-poke visits are defined as visits with *n*_*i*_ = 1 and *ℓ*_*i*_ = 1.

### Binned counts, activity, and group summaries

For a bin width Δ in hours, an event belongs to bin *b* when

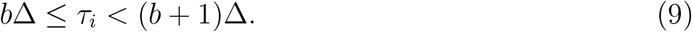

For any event-level value *x*_*i*_, the mouse-level binned count is

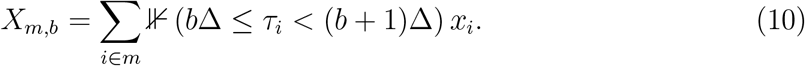

For all-visit activity, *x*_*i*_ = 1 and 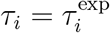. For nose-poke counts, *x*_*i*_ is the number of nose-poke events associated with the visit. For licking counts, *x*_*i*_ is the lick count. For NPA visits and drinking visits, 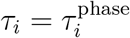 within phase 2, and *x*_*i*_ = 1 for all visits or for lick-positive nose-poke visits, respectively.

Group-level summaries are computed from mouse-level values. For group *g* with *N*_*g*_ contributing mice,

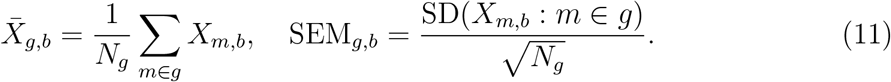

Phase activity summaries first count visits per mouse in hourly phase-relative bins and then report the mean or median visits per hour per mouse and phase. Missing hourly bins within the observed phase window are filled with zero to represent inactive recorded time. These definitions keep plotted activity quantities traceable to mouse-level tables, addressing design goals 3 and 4 in Box 1.

### Bottle consumption and preference

Bottle analyses are computed from nose-poke/lick events with side labels. The user defines which IntelliCage side IDs belong to the left and right bottle sets, allowing the same method to compute total uptake, side-specific uptake, or preference ratios depending on the experiment. This supports the explicit and modular metric design described in goals 3 and 5 of Box 1. For mouse *m* and bin *b*, let

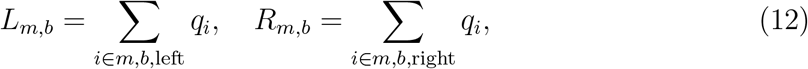

where *q*_*i*_ is the selected consumption column, by default the lick count. Total liquid uptake is

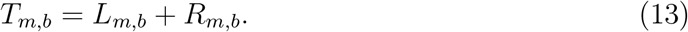

The `calc=“all”` option reports *T*_*m,b*_. The `left_bottle` and `right_bottle` options report *L*_*m,b*_ or *R*_*m,b*_. Ratio options report a side-specific fraction of total consumption:

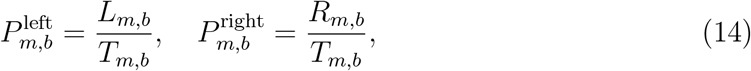

with undefined ratios set to missing when *T*_*m,b*_ = 0.

Optional bin-wise group statistics test whether groups differ in each bin. For two groups, Shapiro tests are used as a normality screen; normally distributed data are compared with Welch’s t-test and non-normal data with the Mann–Whitney U test. For more than two groups, normally distributed data are compared with one-way ANOVA and non-normal data with the Kruskal–Wallis test. Significance stars are reported at *p <* 0.05, *p <* 0.01, and *p <* 0.001.

### Place-learning rates, error rates, and endpoints

For phase *p*, success metric *S*_*i*_, mouse *m*, and segment *d*, the success rate is

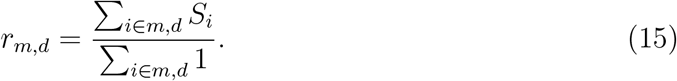

The default metric used in the demonstration is rewarded correct-corner visiting, *S*_*i*_ = *RC*_*i*_. Alternative metrics include *C*_*i*_ for all correct-corner visits and *CN*_*i*_ for correct-corner visits with nose pokes. Error rates use the same denominator and are computed as

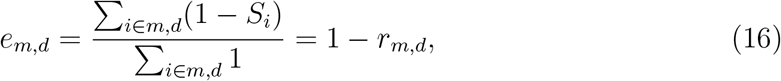

when at least one visit is present in the segment.

Awake/sleep segmentation uses the configured mouse day. If a phase starts at clock hour *h*_*p*_, the package aligns phase elapsed time to the mouse day by

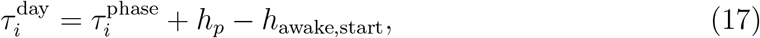

then assigns alternating awake and sleep segments. Endpoint plots collapse a selected phase day and dayphase per mouse, then apply the same rate or error-rate definition. This keeps PL/PR readouts reusable across related protocols while preserving explicit success definitions, as required by design goals 3 and 5 in Box 1.

### Learning onset by experience and time thresholds

Experience-based learning curves are computed over visit number rather than elapsed time, separating self-paced behavioral experience from clock-time phase alignment. This follows design goals 1 and 3 in Box 1 by making the onset criterion explicit while keeping it tied to biological phase windows. For each mouse, visits in a phase are sorted by timestamp. With rolling window size *W*, the success probability for a window starting at visit index *j* is

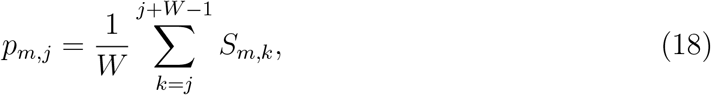

provided the window contains at least the configured minimum number of visits. The window is plotted at center visit *j* + *W/*2. Learning onset is the first center visit of the first run of *K* consecutive windows for which

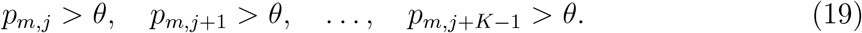

The demonstration used the package defaults of *W* = 20 visits, *K* = 3 consecutive windows, and threshold *θ* = 0.40 for experience-onset analyses unless changed in the user script. Mice that never cross the criterion are retained in the output table with missing onset values and are not forced to an artificial maximum.

The package also supports threshold-onset analyses on clock-time bins. For a binned rate *r*_*m,b*_ and threshold *θ*, the onset is the first non-empty bin start *b*Δ for which *r*_*m,b*_ *> θ*. This definition asks when a mouse first exceeded a predefined binned rate, whereas the experience onset asks after how many self-initiated visits a mouse showed sustained above-threshold performance.

### Derived ratios and reversal components

Derived ratios implement design goal 3 in Box 1 by making composite endpoints traceable to explicitly defined event classes. They count two visit classes within the same mouse, phase day, and dayphase window:

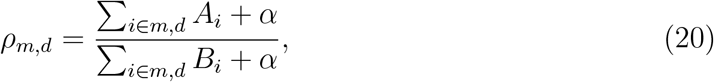

where *A*_*i*_ is the event class of interest, *B*_*i*_ is the reference event class, and *α* is an optional pseudocount. With *α* = 0, ratios with zero denominator are set to missing. Completion efficiency used

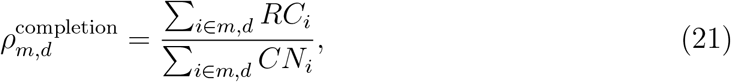

asking how often a correct-corner nose poke was followed by licking. The PR reversal preference index used

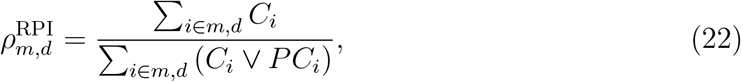

asking, among visits to the two historically relevant corners, what fraction went to the new PR target rather than the previous PL target.

Reversal-component plots classify every PR visit as new correct, previous correct, neutral incorrect corner 1, or neutral incorrect corner 2. For role *q*, the role-specific rate in bin *b* is

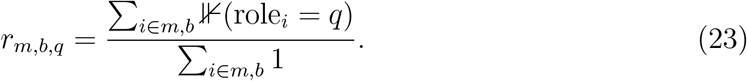

Group summaries are computed as means and SEMs across mice.

### Statistical testing

The package writes test tables for plots that include inferential annotations, so that statistical labels in figures can be inspected through the corresponding output files. This extends the explicit-metric and figure/table-coupling principles in design goals 3 and 4 of Box 1 to default statistical summaries. For two-group endpoint and onset comparisons, the default path evaluates normality and variance structure before choosing a parametric or non-parametric test. When groups satisfy the normality screen and variance assumptions, independent-sample t-tests are used. Otherwise, Mann–Whitney U tests are used. For more than two groups, ANOVA with post hoc Tukey HSD or Kruskal–Wallis with false-discovery-rate-adjusted pairwise Mann–Whitney tests are used depending on the same distributional screening. Chance-level tests for correct-corner rates use binomial tests against the configured chance probability. The statistical outputs are intended as transparent defaults for exploration and reporting, not as a replacement for custom mixed models when the experimental design requires them.

### Synthetic dataset

The synthetic dataset contains two groups of ten mice each and was used as a reproducible test case for the phase-alignment and output-traceability goals in Box 1. Group A was simulated with stronger saccharin engagement, faster acquisition, and stronger reversal learning. Group B was simulated with weaker drinking engagement, slower learning, stronger perseveration at the previously correct corner, and a more difficult reversal phenotype. To test time alignment, Group B was generated as a separate cage run beginning 12 days after Group A and with a 7.5 h delay before the first phase-1 events. Biological phase windows were nevertheless defined in subject metadata, allowing analysis by experimental phase rather than by export-block folder names or absolute calendar time.

The four phases were scheduled as follows: phase 1, free habituation, 0–74 h; phase 2, NPA, 74–122 h; phase 3, PL, 122–194 h; and phase 4, PR, 194–266 h. PL and PR were limited to 72 h for common-window analyses. The demonstration is synthetic and did not involve animal experimentation.

### Software and output generation

The toolkit is implemented in Python and uses the scientific Python ecosystem for tabular analysis, statistics, and plotting. Each analysis run writes plot files together with TSV/CSV-compatible tables, YAML settings, loaded metadata, phase manifests, and audit-oriented intermediate outputs. This makes figures traceable to machine-readable data and makes it possible to rerun an analysis from the same user script and raw data layout, addressing the reproducible-output and FAIR-oriented sharing goals in Box 1.

## Data availability

The synthetic IntelliCage example-data archive used for the analyses reported here is publicly available via Zenodo [93]. It includes the synthetic Group A/B place-learning/place-reversal dataset, IntelliCage-style raw export tables, metadata, manifests, and generated result files needed to inspect the demonstration workflow.

## Code availability

*ic-analysis* is distributed under the GNU General Public License v3.0 or later. The archived software release is publicly available via Zenodo [94]. The source code, documentation, issue tracker, and example user scripts are hosted in the public repository at https://github.com/FabrizioMusacchio/ic-analysis. The repository contains the analysis scripts and synthetic-data workflow used to reproduce the demonstration analyses reported here.

## Acknowledgments

We thank the open-source community for developing and maintaining the scientific Python tools on which *ic-analysis* relies, including NumPy [95], pandas [96], SciPy [97], Matplotlib [98], statsmodels [99], and PyYAML [100].

## Funding

M.F. received funding from the European Research Council, ERC MicroSynCom 865618, and the German Research Foundation, DFG SFB1089 C01/B06 and SPP2395.

## Use of AI-assisted tools

The authors used OpenAI ChatGPT, GPT-5, to assist with language editing and wording refinement. The tools were not used to generate primary research data, perform unsupervised interpretation of results, or make authorship-level scientific decisions. All AI-assisted text changes were reviewed, verified, and edited by the authors, who take full responsibility for the accuracy, integrity, and content of the submitted article.

## Author contributions

F.M. conceptualized the study, developed the software, designed and generated the synthetic dataset, performed the analyses, created the figures, curated the project resources, and wrote the manuscript. M.F. supervised the project, acquired funding, and reviewed and edited the manuscript.

## Notes

### Competing Interest Statement

The authors have declared no competing interest.

